# Single-cell foundation models transfer across deep evolutionary distances

**DOI:** 10.64898/2026.10.28.762457

**Authors:** Lena Trnovec, Mariko Katoh-Kurasawa, Rachel N. Arey, Gad Shaulsky, Blaž Zupan

## Abstract

Single-cell foundation models are trained mostly on human and other vertebrate cells, yet are proposed as general-purpose models. Whether they transfer far beyond their training species is unknown. We evaluated zero-shot embeddings from Geneformer, scGPT and UCE against PCA and scVI in developmental atlases of *Dictyostelium discoideum* and *Caenorhabditis elegans*, separated from humans by 1.2 billion and 600 million years. Without retraining, the models recovered developmental order and preserved organism-specific cell identities. UCE encodes genes by protein sequence and could use 97% of measured genes, whereas Geneformer and scGPT, after ortholog mapping, used under one-fifth. UCE performed most consistently, often exceeding species-native baselines. Restricted to ortholog-mappable genes, it still scored above vocabulary-based models on five of six annotation tasks, some narrowly. Foundation models can thus yield informative cell embeddings across more than a billion years of evolution, and protein-sequence gene representations make these embeddings more complete and often more accurate.

## Introduction

Single-cell RNA sequencing has enabled atlas-scale descriptions of cell identity, state and developmental progression across tissues and organisms ^1–3^. In parallel, large self-supervised models have been proposed as reusable representations for single-cell transcriptomics ^4–11^. Their appeal is broad transfer: an embedding learned from many cells could support a new biological analysis without training a task-specific model. Existing evaluations, however, show that performance depends on the dataset, endpoint, preprocessing and baseline, and that pretrained models do not reliably outperform simpler representations ^12–17^.

Applying these models to evolutionarily distant species reveals a constraint that is less obvious in human-centered benchmarks: a model can process only genes that are recognized by its input representation, but many single-cell foundation models use a fixed human gene vocabulary. Applying such a model to another species therefore requires a preprocessing workflow that is external to the model: source genes must be linked to human orthologs, ambiguous mappings must be resolved, and genes without an accepted target must be discarded. Orthology is not a simple lookup, particularly after gene duplication, loss and functional divergence ^18–20^. Different resources and confidence rules can consequently expose a model to different subsets of the same transcriptome. Model performance and mapping coverage are inseparable unless both are reported.

Cross-species methods handle the issue of ortholog mapping in different ways. SAMap, CAME and SATURN incorporate homology graphs or protein-derived gene representations to align cells across species^21–23^. Broader benchmarks find that phylogenetic distance and gene representation shape integration performance^24,25^. Universal Cell Embedding (UCE) takes a related representation-level approach: each gene is associated with an ESM-2 protein embedding, allowing the model input to be constructed from the target species’ proteome rather than from a human gene symbol alone ^11,26,27^. This approach improves coverage, but it does not guarantee that a protein representation trained from sequence will preserve every evolutionary or cellular relationship needed downstream. Recent work distinguishes successful sequence modeling from explicit evolutionary reasoning, cautioning against treating geometric proximity in protein-embedding space as a complete biological mechanism ^28^.

We studied this issue in a *zero-shot cross-species* setting: pretrained models were applied to the target atlas without retraining, fine-tuning, label supervision or cross-species cell alignment. The embedding may still require model-specific input construction, including ortholog mapping for fixed-vocabulary models ^6,7,11^. This distinction matters operationally. A method that requires a multi-resource mapping pipeline has a larger deployment burden than one that can use target-species protein sequences, even when both ultimately produce embeddings without updating model parameters. Reducing this burden would also make advanced representation methods more accessible for non-model organisms whose data are poorly served by human-centered vocabularies.

Developmental atlases provide two complementary evaluation axes. Their cells carry discrete biological annotations, and they also lie along an external temporal process, permitting evaluation beyond visual cluster separation. We selected two organisms that are evolutionarily distant from humans and from each other. *Dictyostelium discoideum*, a social amoeba separated from humans by approximately 1.2 billion years ^29^, develops when starved cells aggregate into a multicellular structure with prespore and prestalk fates ^30–33^. *Caenorhabditis elegans*, separated from humans by approximately 600 million years ^29^, follows an invariant metazoan lineage with parallel tissue specification^34,35^. The *D. discoideum* atlas has direct collection times across development, whereas the *C. elegans* atlas uses embryo time estimated by correlation with a whole-embryo RNA-seq series^35,36^. The two systems therefore challenge representations through different gene repertoires, developmental strategies and temporal labels. Because our goal was to evaluate behavior under substantial evolutionary divergence, we prioritized organismal distance and developmental coverage over the number of included datasets.

We hypothesized that, for distant species, the transferability of a single-cell foundation model is governed less by the species it was trained on than by how it represents genes: a fixed human vocabulary restricts the model to the conserved, mappable fraction of the target transcriptome, whereas a protein-sequence representation can expose nearly all protein-coding genes. To test this, we compared five methods spanning three representation strategies: PCA^37^and scVI^38^use the full species-native expression matrix; Geneformer ^6^and scGPT^7^require conversion to a human ortholog vocabulary; and UCE uses protein-derived gene embeddings ^11^. We asked whether models pretrained on vertebrate cells yield biologically meaningful zero-shot embeddings for organisms up to 1.2 billion years distant; how many measured genes each method can use after all vocabulary filters; how well each embedding recovers developmental order and curated organism-specific annotations; and, by restricting UCE-4L to the Tier 1–2 ortholog-available gene pool, how much of any difference reflects gene access rather than representation.

The study makes three contributions. First, it extends zero-shot evaluation of single-cell foundation models beyond vertebrates, to a nematode and to a social amoeba outside the animal kingdom. We show that useful transfer is possible at these distances. Second, it identifies gene accessibility as a distinct and quantifiable dimension of cross-species performance, and uses coverage-aligned controls to separate it from representation quality. Third, it translates these findings into reporting requirements for cross-species benchmarks. Together, these analyses show that single-cell foundation models trained on vertebrate cells can produce informative cell embeddings for organisms separated from humans by more than a billion years of evolution. How much of the target organism’s biology these embeddings capture depends on how the model represents genes, which determines both how many of the target organism’s genes the model can use and how informatively it encodes them.

## Results

### Two developmental systems provide complementary cross-species tests

The *D. discoideum* atlas ^33^ contained 48,983 cells and 12,883 genes sampled in replicate at 4-hour intervals across the first 20 hours of starvation-induced development (Fig. 1a–c). The series spanned starvation, aggregation, mound and slug formation, and culmination. Developmental time was therefore a directly observed collection label. The processed *C. elegans* matrix was derived from a published atlas ^35^ and contained 89,701 embryonic cell profiles and 20,222 genes, with founder-lineage structure and progressive tissue diversification (Fig. 2a–c). Of these profiles, 86,024 cells were marked as passing the source study’s initial quality control or later whitelist, matching its reported final dataset, and 3,677 cells were marked otherwise; all were retained in the benchmark. The embryo-time coordinate was inferred by correlation with a whole-embryo RNA-seq time series rather than measured as a collection time ^36^. Agreement with that coordinate tested recovery of a published developmental estimate, not recovery of an independent experimental timestamp.

**Figure 1:**
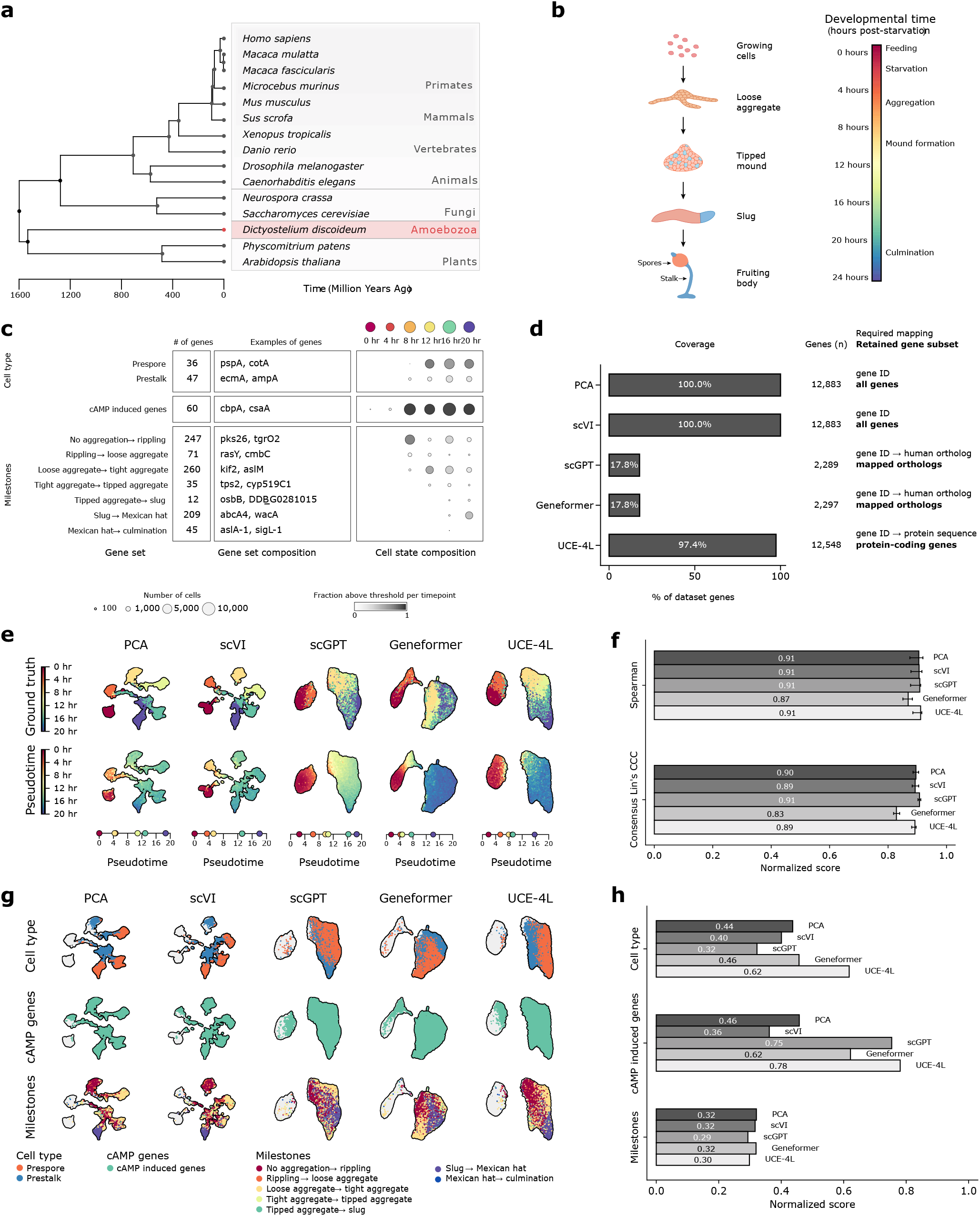
Cross-species benchmarking in the social amoeba *D. discoideum*. **(a)** TimeTree-based timetree showing estimated evolutionary distances from *Homo sapiens* (human); horizontal branch lengths and the axis denote millions of years before present, whereas vertical order is graphical. Orange taxa are the eight species represented in the UCE training corpus (*H. sapiens, Macaca mulatta, Macaca fascicularis, Microcebus murinus, Mus musculus, Sus scrofa, Xenopus tropicalis* and *Danio rerio*). Additional reference taxa include the benchmark species *Caenorhabditis elegans* and *Dictyostelium discoideum*. **(b)** Developmental schematic spanning growth, starvation, aggregation, mound formation, slug and culmination, with corresponding sampling times after starvation. **(c)** Dataset overview of three curated annotation categories: prespore versus prestalk identity, cAMP-pulse-induced genes and developmental milestones. The table gives set size and representative genes; the bubble plot shows the fraction of cells above the activity threshold at each timepoint, with bubble area proportional to cell count. **(d)** Percentage and absolute number of genes retained by each method; the required mapping or protein-asset filter and final subset are printed in the panel. **(e)** UMAPs colored by observed collection time (top) and diffusion pseudotime (bottom). Each dot is a cell; UMAP axes are arbitrary and local neighborhoods, rather than axis values, convey proximity. A black underlay outlines the point cloud for legibility and is not an inferred cluster boundary. The one-dimensional plot gives mean collection time in six successive pseudotime bins. **(f)** Spearman rank correlation (*ρ*) and consensus Lin’s concordance correlation coefficient (CCC) between pseudotime and collection time, rescaled to [0,1] by (*r*+1)*/*2; higher values indicate greater agreement. Error bars are 95% repeated-subsampling stability intervals from 30 paired repeats. **(g)** UMAPs colored by the three annotation categories in **c**: prespore versus prestalk identity, cAMP-pulse-induced genes and developmental milestones. **(h)** avgBIO, the mean of NMI, ARI and label ASW, for each annotation category; higher values indicate greater label preservation.

**Figure 2:**
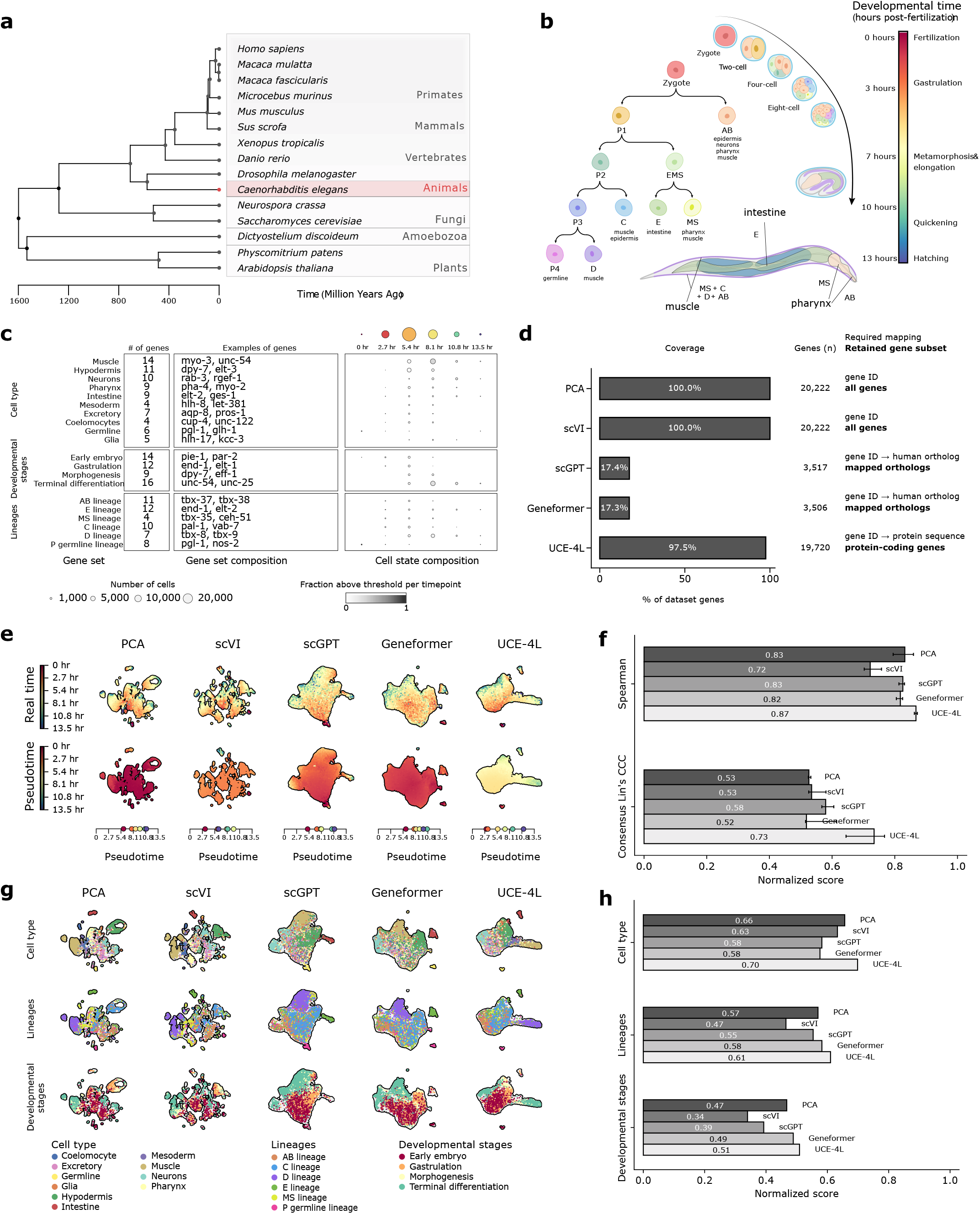
Cross-species benchmarking in the nematode *C. elegans*. **(a)** TimeTree-based timetree showing estimated evolutionary distances from humans; horizontal branch lengths and the axis denote millions of years before present, whereas vertical order is graphical. Orange taxa are the eight species represented in the UCE training corpus; the remaining taxa provide evolutionary context and include both benchmark organisms. **(b)** Embryonic developmental schematic with founder-cell lineages (AB, E, MS, C, D, germline), major tissue outputs and approximate developmental times. **(c)** Dataset overview of three curated annotation categories: cell/tissue classes, lineage categories and developmental markers. The table gives set size and representative genes; the bubble plot shows the fraction of cells above the activity threshold across embryo-time bins, with bubble area proportional to cell count. **(d)** Percentage and absolute number of genes retained by each method; the required mapping or protein-asset filter and final subset are printed in the panel. **(e)** UMAPs colored by the published embryo-time estimate (top) and diffusion pseudotime (bottom). Each dot is a cell; UMAP axes are arbitrary and local neighborhoods convey proximity. A black underlay outlines the point cloud for legibility and is not an inferred cluster boundary. The one-dimensional plot gives mean estimated embryo time in six successive pseudotime bins. **(f)** Spearman rank correlation (*ρ*) and consensus Lin’s CCC between pseudotime and estimated embryo time, rescaled to [0,1] by (*r*+1)*/*2; higher values indicate greater agreement. Error bars are 95% repeated-subsampling stability intervals from 30 paired repeats. **(g)** UMAPs colored by the three annotation categories in **c**: cell/tissue classes, lineage categories and developmental markers. **(h)** avgBIO, the mean of NMI, ARI and label ASW, for each annotation category; higher values indicate greater label preservation.

For each organism, we constructed three annotation categories from published marker sets (Fig. 1c; Fig. 2c). Complete marker-gene lists, set memberships and panel-level literature references are provided in Supplementary Data 1. The *D. discoideum* categories describe prespore versus prestalk identity, grounded in in situ RNA hybridization and RNA-seq of biophysically separated cells^39,40^; genes induced by cAMP-pulse signaling during early development^41–43^; and milestone genes up-regulated at defined developmental transitions ^41,44,45^. The *C. elegans* categories describe cell/tissue classes, lineage categories across the AB, E, MS, C, D and germline branches, and developmental markers^34,35^. We used these expression-derived labels to evaluate how well each method preserved the corresponding biological annotations. They are not independent experimental outcomes.

### Human-vocabulary models see less than one-fifth of distant transcriptomes

We first counted genes after applying all the method-specific input filters (Fig. 1d; Fig. 2d). PCA was computed and scVI was trained separately on the full species-native expression matrices—12,883 genes for *D. discoideum* and 20,222 for *C. elegans*—because neither required a pretrained human vocabulary. The stored Tier 1–2 ortholog annotations used to generate the recorded embeddings selected 2,337 source-gene features in *D. discoideum* and 3,568 in *C. elegans* (Supplementary Table S1). Mapping these features to model identifiers, excluding identifiers absent from each fixed human vocabulary and collapsing duplicate target identifiers left 2,297 genes (17.8%) for Geneformer and 2,289 (17.8%) for scGPT in *D. discoideum*. In *C. elegans*, the same final filters left 3,506 genes (17.3%) for Geneformer and 3,517 (17.4%) for scGPT.

UCE could use 12,548 genes (97.4%) in *D. discoideum* and 19,720 genes (97.5%) in *C. elegans*. Its filter requires a protein embedding and chromosome-position input recognized by the UCE pipeline, but not a one-to-one human gene symbol. Genes without an annotated protein product, including many non-protein-coding genes, remain unavailable^11,23^. Thus UCE also performs input selection; the difference is that the selection is based primarily on the target-species protein assets rather than on a human ortholog vocabulary.

Coverage was both a scientific and an operational result. The ortholog pipeline combined OrthoFinder, OrthoDB, OMA and curated organism resources before applying confidence rules and model-vocabulary filters (Supplementary Fig. S1). Even after this multi-source procedure, more than four-fifths of the measured genes were absent from Geneformer and scGPT inputs. The excluded set can contain lineage-specific genes, paralogs and rapidly diverging genes, but coverage alone does not establish that every excluded gene is informative. The loss nevertheless extended to genes of known biological relevance: of 83 curated *D. discoideum* cell-type markers, only 2 survived Tier 1–2 ortholog mapping, whereas all 83 were eligible for UCE (Supplementary Table S2). We therefore treated the observed coverage as a property of these organisms, mapping rules and model releases rather than as a universal percentage for each method.

### Foundation models do not uniformly improve developmental ordering

To test whether differences in gene availability accompanied differences in model performance, we first evaluated developmental progression by computing diffusion pseudotime (DPT)^46^ separately in each representation. In the UMAPs, each point is a cell, color denotes the reference developmental coordinate or inferred pseudotime, and proximity represents neighborhood structure in a two-dimensional projection. The *D. discoideum* maps broadly separated early single-cell stages from later multicellular stages, whereas the *C. elegans* maps contained several branches consistent with parallel lineage specification. The degree of fragmentation and local mixing differed among the methods, motivating quantitative comparison with root-averaged Spearman correlation and consensus Lin’s concordance correlation coefficient (CCC)^47^. These values are shown in Fig. 1f and Fig. 2f, and the full values are provided in Table 1.

**Table 1:** Summary of benchmark performance across methods and organisms. The best value for each metric or annotation is shown in bold.

| Method | *D. discoideum* |  |  |  | *C. elegans* |  |  |  |
|---|---|---|---|---|---|---|---|---|
|  | Pseudotime | Cell type | cAMP-pulse-induced genes | Milestones | Pseudotime | Cell/tissue classes | Lineage categories | Developmental markers |
| PCA | .905 (.895) | .437 | .458 | **.321** | .832 (.526) | .657 | .569 | .468 |
| scVI | .906 (.893) | .401 | .362 | .316 | .722 (.535) | .633 | .465 | .340 |
| Geneformer | .869 (.829) | .457 | .622 | .319 | .818 (.518) | .576 | .582 | .489 |
| scGPT | .909 (**.908**) | .322 | .754 | .293 | .826 (.579) | .582 | .554 | .393 |
| UCE-4L | **.911** (.892) | **.618** | **.781** | .298 | **.868** (**.734**) | **.699** | **.610** | **.509** |

*Note.* Pseudotime entries give root-averaged Spearman *ρ* with consensus Lin's CCC in parentheses; both are rescaled to [0, 1] by (*r* + 1)/2. Annotation columns report avgBIO, the unweighted mean of NMI, ARI and label ASW.

In *D. discoideum*, all methods recovered the developmental ordering (rescaled Spearman *ρ* = 0.869– 0.911; Table 1). UCE-4L had the highest point estimate for rank agreement (0.911), whereas scGPT had the highest consensus concordance (0.908). PCA (0.905 rank agreement), scVI (0.906) and scGPT (0.909) were closely grouped with UCE-4L. Paired stability intervals did not separate UCE-4L from PCA, scVI or scGPT for rank agreement; Geneformer was lower. The narrow leading range is consistent with a strong temporal signal in the *D. discoideum* transcriptome ^41,45^, which does not require a pretrained representation to recover.

In *C. elegans* (Fig. 2f), UCE-4L achieved the highest rescaled rank agreement (0.868), with a robust advantage over every comparator in the paired repeated-subsampling analysis. UCE-4L also had the highest rescaled consensus concordance (0.734), followed by scGPT (0.579). Consensus concordance was lower than rank agreement for every method. This pattern is compatible with several features of the endpoint: embryogenesis contains parallel lineage trajectories rather than a single path, and the reference coordinate is itself an inferred embryo-time estimate ^36^. It should not be interpreted as an absolute timing benchmark.

Together, the two datasets showed that pseudotime performance was context dependent. A pretrained representation offered little separation from strong baselines when the temporal signal was dominant, whereas UCE-4L improved both rank agreement and consensus concordance in the more heterogeneous embryo. The differing margins across metrics and organisms argue against collapsing trajectory quality to a single context-free ranking.

Because the UMAPs and binned plots are reduced summaries, we based cross-method comparisons on cell-level correlations across each full atlas and used repeated subsampling to assess their stability (Methods; Supplementary Table S3).

### Model performance varies across annotation categories

We next tested whether each model retained the curated labels as clusterable and geometrically separated structures. In the *D. discoideum* annotation-colored UMAPs, UCE-4L placed prespore and prestalk cells in adjacent but visibly distinct regions, whereas their separation was more diffuse or fragmented in several other representations. Cells positive for cAMP-pulse-induced genes occupied localized regions in the scGPT, Geneformer and UCE-4L maps, consistent with their higher scores than PCA and scVI. By contrast, the milestone labels remained intermingled along a developmental continuum in every representation, matching the narrow range of lower avgBIO values (Fig. 1g,h and Table 1).

In *C. elegans*, cell/tissue classes occupied broad regions in every map and were most coherently organized in the UCE-4L, PCA and scVI representations. Lineage categories remained partially over-lapping but showed more localized regions in UCE-4L and Geneformer than in several comparators.

Developmental-stage labels overlapped most extensively, consistent with their lower scores and their graded temporal structure (Fig. 2g,h and Table 1).

We quantified these qualitative patterns with avgBIO, the unweighted mean of normalized mutual information (NMI), adjusted Rand index (ARI) and label average silhouette width (ASW)^7,48–50^; the component values are reported in Supplementary Table S4. Because Leiden resolution was optimized against each label and because all cells came from one atlas per species, these scores are descriptive within-dataset summaries rather than estimates of out-of-sample classification performance.

In *D. discoideum*, UCE-4L achieved the highest avgBIO for cell type (0.618), ahead of the next-best method, Geneformer (0.457), and for cAMP-pulse-induced genes (0.781), narrowly above scGPT (0.754; Supplementary Table S4). Milestones were harder for every method (all avgBIO 0.321), with no method standing out (PCA highest at *≤* 0.321; UCE-4L 0.298). In *C. elegans*, UCE-4L gave the highest avgBIO on all three panels—cell/tissue classes (0.699), lineage categories (0.610) and developmental markers (0.509)—though by small margins over the next-best method in each case. PCA was next for cell/tissue classes (0.657), while Geneformer was next for lineage categories (0.582) and developmental markers (0.489; Supplementary Table S4). Across the six headline panels UCE-4L ranked first on five. Its clearest difference was for *D. discoideum* cell type (0.618 versus 0.457 for Geneformer); leading differences on the other panels were smaller. We therefore described UCE-4L as the most consistent method in the headline comparison in this benchmark, not as uniformly or significantly superior.

Following the reporting convention used in scIB-style comparisons^51^, no within-dataset confidence intervals or formal tests were attached to avgBIO point estimates.

### Gene coverage explains some, but not all, of the protein-based advantage

The main comparison changes both gene set and representation. Full-coverage UCE-4L uses nearly all protein-coding genes, whereas Geneformer and scGPT use a much smaller human-ortholog subset. We therefore recomputed UCE-4L on the Tier 1–2 ortholog-available source-gene pool used before the human-vocabulary filters (Supplementary Table S5). This was a coverage-aligned control within UCE, not a complete architectural control; the final usable identifiers, tokenization, model training and aggregation still differ between methods.

For cAMP-pulse-induced genes in *D. discoideum*, restricting UCE-4L to ortholog-available genes had almost no effect (avgBIO 0.770 versus 0.781 for full-coverage UCE-4L), and the ortholog-restricted UCE-4L condition remained above Geneformer (0.622) and scGPT (0.754). For *D. discoideum* milestones, where full-coverage UCE-4L did not lead (0.298), ortholog-restricted UCE-4L scored 0.332, slightly above Geneformer (0.319) and scGPT (0.293); all methods remained closely grouped on this panel. In *C. elegans*, ortholog-restricted UCE-4L stayed close to full-coverage UCE-4L for cell/tissue classes (0.655 versus 0.699), lineage categories (0.593 versus 0.610) and developmental markers (0.491 versus 0.509), and remained above both ortholog-vocabulary models on each panel. Gene count alone therefore does not explain those differences. The result is consistent with a contribution from protein-based encoding, while remaining observational because representation and architecture were not independently randomized.

By contrast, ortholog-restricted UCE-4L dropped from 0.618 to 0.342 for *D. discoideum* cell type and fell below Geneformer (0.457). Marker retention explains this contrast: only 2% of *D. discoideum* cell-type markers survived Tier 1–2 ortholog mapping, compared with 21–28% of the markers in each *C. elegans* panel (Supplementary Table S2). In this panel, access to additional target-species genes is therefore a more plausible explanation than encoding alone. The controls therefore suggest task-dependent contributions from coverage and representation.

The same restriction did not reduce rank-based developmental ordering (Supplementary Table S6). In *D. discoideum*, rank agreement was 0.914 for ortholog-restricted UCE-4L versus 0.911 for full-coverage UCE-4L, and consensus concordance was 0.903 versus 0.892. In *C. elegans*, rank agreement was unchanged after rounding (0.868 versus 0.868), while consensus concordance decreased from 0.734 to 0.708 but remained above the headline comparators. Coverage therefore affected the *D. discoideum* cell-type annotation strongly without accounting for the UCE-4L rank-ordering result in either atlas.

### Conventional preprocessing changes several PCA comparisons

Because the headline PCA condition was deliberately minimal and baseline construction can influence benchmark conclusions, we tested how a conventional single-cell PCA recipe changed its performance. After selection of 2,000 highly variable genes, library-size normalization, log1p transformation and scaling, *D. discoideum* avgBIO increased from 0.437 to 0.531 for cell type, from 0.458 to 0.484 for cAMP-pulse-induced genes and from 0.321 to 0.387 for milestones. In *C. elegans*, it increased from 0.657 to 0.724 for cell/tissue classes and from 0.569 to 0.643 for lineage categories, but decreased from 0.468 to 0.387 for developmental markers (Supplementary Table S7). Processed PCA therefore surpassed UCE-4L on three panels: *D. discoideum* milestones and *C. elegans* cell/tissue classes and lineage categories. It remained below UCE-4L for *D. discoideum* cell type and cAMP-pulse-induced genes and for *C. elegans* developmental markers.

The same preprocessing did not uniformly improve trajectory recovery. In *D. discoideum*, rescaled rank agreement changed from 0.905 to 0.881 and consensus concordance from 0.895 to 0.884. In *C. elegans*, rank agreement decreased from 0.832 to 0.722, while consensus concordance remained similar (0.526 versus 0.525; Supplementary Table S8). Thus PCA performance depended on both its recipe and the endpoint: conventional preprocessing improved five of six annotation comparisons but weakened rank-based developmental ordering in both organisms, and UCE-4L’s five-of-six annotation ranking was not invariant to reasonable baseline construction.

### A common output dimension preserves the main annotation pattern

The headline representations also differed in dimensionality: PCA and scVI were 100-dimensional, Geneformer and scGPT were 512-dimensional, and UCE-4L was 1,280-dimensional. To test whether the larger representations benefited simply from additional coordinates, we retained PCA and scVI unchanged and reduced the other three representations independently to 100 principal components (Supplementary Table S9). The projected representations retained 95.1–98.1% of their variance across methods and organisms.

UCE-4L remained highest for *D. discoideum* cell type (0.607) and cAMP-pulse-induced genes (0.782), and for *C. elegans* cell and tissue classes (0.698), developmental markers (0.510) and lineage categories (0.612). These values were close to the corresponding native-dimensional scores (0.618, 0.781, 0.699, 0.509 and 0.610). For *D. discoideum* milestones, projected Geneformer and unchanged PCA both rounded to 0.321, and all methods remained closely grouped. Thus unequal output dimensionality did not account for the clear annotation differences, although post-embedding PCA changed individual scores and was not an architectural control.

UCE-4L developmental ordering was also stable to this reduction (Supplementary Table S10). Its rank agreement was 0.912 at 100 dimensions versus 0.911 natively in *D. discoideum*, and 0.866 versus 0.868 in *C. elegans*; the corresponding consensus-concordance values were 0.892 versus 0.892 and 0.739 versus 0.734. The largest change among the other developmental-ordering comparisons was an increase in *C. elegans* Geneformer consensus concordance from 0.518 to 0.586, which altered the ordering of the middle-ranked methods but not the leading method. Thus the common-dimensionality analysis does not explain the UCE-4L trajectory result, although post-embedding reduction can affect individual metrics.

### UCE checkpoint depth is organism dependent

UCE was released with 4- and 33-layer cell-transformer checkpoints, so we tested whether the comparison depended on checkpoint depth (Supplementary Table S11). In *D. discoideum*, UCE-33L had lower avgBIO than UCE-4L on all three annotation panels and lower rank agreement, but slightly higher consensus concordance (0.904 versus 0.892). In *C. elegans*, the two checkpoints had similar annotation scores, while UCE-4L had both higher rank agreement and higher consensus concordance (0.734 versus 0.646).

We next examined whether protein-space distribution shift could contribute to this organism-specific depth effect (Supplementary Table S12). Before calibration, mean nearest-human cosine was lower for *D. discoideum* than for *C. elegans* protein vectors (0.761 versus 0.813). Global moment matching and a macrogene-residual adapter increased the *D. discoideum* value to 0.806 and 0.809, respectively, and partially rescued UCE-33L avgBIO for cell type (0.252 to 0.430 and 0.591) and cAMP-pulse-induced genes (0.500 to 0.771 and 0.769). The adapters did not improve every condition: global matching reduced UCE-4L *D. discoideum* cell-type avgBIO from 0.618 to 0.456, and neither adapter consistently improved UCE-33L in *C. elegans*. We retained UCE-4L for the headline comparison because it was more consistent across the two organisms. These exploratory analyses do not establish that shallower checkpoints are generally preferable or that protein-space shift caused the depth effect.

## Discussion

Single-cell foundation models can transfer surprisingly far across evolution. Applied without retraining to a nematode and to a social amoeba separated from humans by about 600 million and 1.2 billion years, respectively, models pretrained on vertebrate cells recovered developmental order and preserved organism-specific cell identities, lineages and signaling states. This was far from guaranteed, as neither target organism nor any close relative was represented in the single-cell training data. The limitation is therefore not simply whether transfer is possible, but how gene representation determines how much of the target organism a model can see, and therefore how well transfer works. A fixed human vocabulary discarded more than four-fifths of the measured genes before inference, even after a multi-resource ortholog workflow, whereas protein-based input retained nearly all protein-coding genes. Part of the performance difference thus originates upstream of the model, and benchmarks that report only downstream scores confound representation quality with input accessibility.

The ortholog-pool restriction helps distinguish two components of that result. On *D. discoideum* cell type, restricting UCE-4L to ortholog-available genes removed most of its advantage, consistent with coverage being central for a marker collection poorly retained by ortholog mapping. On cAMP-pulse-induced genes and all three *C. elegans* panels, much of the UCE-4L score persisted after the restriction, and ortholog-restricted UCE-4L remained above both ortholog-vocabulary models. Gene count alone is therefore insufficient to explain the UCE-4L advantage on these four panels, implicating properties of the protein-based representation or the downstream architecture. The sixth panel, *D. discoideum* milestones, offers no such test because full-coverage UCE-4L did not lead there; ortholog-restricted UCE-4L scored slightly above both ortholog-vocabulary models, but all methods were closely grouped. This interpretation is compatible with UCE and SATURN, which use protein language models to place genes into a shared representation ^11,23^. It is not a causal decomposition: the final model-usable identifiers differ slightly even after the shared source-gene restriction, and UCE, Geneformer and scGPT also differ in architecture, training data, tokenization and cell-level aggregation.

The checkpoint comparison reinforces the importance of reporting model configuration. UCE-33L was substantially weaker than UCE-4L in *D. discoideum* but similar in *C. elegans*. Lower nearest-human protein similarity in *D. discoideum* and partial rescue after label-free alignment suggest that protein-space shift contributes to this sensitivity, although calibration did not improve every endpoint and the analysis does not establish shift as the sole cause.

Strong species-native baselines remain essential. PCA and scVI used all the measured genes without pretrained biological priors. They recovered the dominant *D. discoideum* temporal axis almost as well as the leading pretrained models, and PCA was competitive on several annotation panels. Conventional preprocessing strengthened PCA on five of six annotation panels, enough to exceed UCE-4L on three panels, but weakened rank-based developmental ordering in both organisms. This pattern matches recent single-cell foundation-model evaluations showing that gains can be task dependent and that baseline choice materially changes the conclusion ^15–17^. Evaluations of pretrained embeddings should therefore include a simple, fully specified baseline and test whether reasonable preprocessing choices change the comparison.

The benchmark also exposes a mismatch between annotation structure and common embedding metrics. Fate-localized annotations were easier to recover than graded developmental annotations. NMI and ARI reward a clustering geometry, while label ASW rewards compact separation. A developmental continuum can preserve meaningful order without forming discrete clusters, which helps explain why pseudotime and avgBIO need not identify the same leading method. Reporting both endpoints avoids treating a single composite as a complete measure of biological utility.

These conclusions hold within boundaries set by the study design. First, the benchmark included only two developmental atlases. The design prioritized large developmental atlases from deeply separated lineages with interpretable temporal structure, making it a targeted test of cross-species robustness rather than a survey; it did not represent the diversity of clades, perturbations or sequencing protocols in which cross-species transfer could be used. We also lacked an in-distribution or closely related-species reference, so the design could not estimate how performance or mapping coverage degrades continuously with evolutionary distance.

Second, the benchmark focused on developmental ordering and marker-derived annotation preservation. We selected these complementary endpoints because they could be evaluated directly from the embeddings and therefore isolated zero-shot representational quality for continuous developmental organization and discrete biological annotation. Tasks such as supervised label transfer, perturbation prediction, regulatory-network inference and direct cross-species alignment introduce additional training data, optimization choices and task-specific models that can obscure differences attributable to the embeddings themselves. The study therefore did not evaluate every downstream application or compare the wider range of algorithms designed specifically for these endpoints.

Third, marker labels were constructed from the same expression matrices that were embedded. This enabled organism-specific evaluation when independent experimental labels were unavailable, but it also coupled the endpoint to marker selection, thresholding and annotation quality. Literature-based marker curation, expression-matched nulls and explicit multiple-testing control reduced arbitrariness without making the labels independent outcomes. Fourth, the ortholog consensus combined several resources under explicit tiers because individual resources use different evidence and can disagree after duplication, loss and divergence ^18–20^. The consensus made the mapping procedure reproducible and conservative, but it did not remove orthology uncertainty or the simplification of many-to-many relationships required by one-to-one model vocabularies. Different databases, releases or conflict rules could therefore change coverage.

Fifth, avgBIO values were descriptive full-data point estimates. Optimizing Leiden resolution against the target label made the comparison useful for representation auditing but unsuitable as an unbiased estimate of performance on unseen cells. We therefore did not attach within-dataset significance claims to the avgBIO rankings. Pseudotime intervals instead measured stability under repeated cell subsampling and graph reconstruction; because repeats overlapped, they were not classical population confidence intervals. Finally, the benchmark evaluated specific released configurations rather than exhaustively searching each model. We compared both released UCE depths and disclosed that checkpoint sensitivity influenced the headline choice, but other configurations could change scores.

Within these boundaries, the results described here have a direct consequence for how cross-species foundation models could be evaluated and used. Downstream rankings shifted with reasonable analytical choices, which included baseline preprocessing, checkpoint depth and, for one panel, gene coverage.by each model’s input representation, (iv) the retention of biologically relevant markers, and (v) performance under coverage-aligned controls and against species-native baselines. More broadly, foundation models that serve the diversity of life may not need training data from every organism, but they do need input representations that can access its genes.

## Methods

### Datasets and stored expression matrices

We analyzed processed matrices derived from two published developmental single-cell RNA-seq atlases. The *D. discoideum* matrix^33^ contains 48,983 cells and 12,883 genes sampled in replicate at 4-hour intervals from 0 to 20 hours after starvation. The stored developmental target was the discrete collection-time label. The *C. elegans* matrix derived from Packer et al. ^35^ contains 89,701 embryonic cell profiles and 20,222 genes. The source paper reports a final integrated dataset of 86,024 cells; the local matrix contains those profiles, marked as passing initial quality control or a later whitelist, plus 3,677 profiles marked otherwise. We retained all profiles in the supplied matrix. Its developmental target was the published embryo.time estimate, derived by correlation with a whole-embryo RNA-seq time series ^36^, rather than a direct collection timestamp. Both processed AnnData files retain sparse raw-count matrices in X and a counts layer. We did not remove additional cells or genes for the main benchmark, and no expression-based selection was applied before the full-gene PCA or scVI conditions.

### Benchmark conditions

The headline comparison contains full-gene PCA and scVI, Tier 1–2 Geneformer and scGPT, and the standard pretrained UCE-4L model. Their dimensions were 100, 100, 512, 512 and 1,280. Raw-count PCA was retained as a minimal, full-gene baseline because no equivalent target-dataset preprocessing was inserted before the pretrained embeddings; conventionally preprocessed PCA was evaluated as a sensitivity analysis. UCE-4L was selected after comparison with the released 33-layer checkpoint; complete UCE-33L results are reported in the supplement.

### Embedding generation

The headline PCA baseline was computed separately from each raw-count matrix using all genes and 100 components, without normalization, log transformation, highly-variable-gene selection or scaling. A sensitivity condition used 2,000 highly variable genes selected with the Seurat-v3 method, normalization to 10,000 counts per cell, log1p, scaling capped at 10, and 100 components. No batch correction was applied. scVI^38^ was trained separately on the full count matrices with a negative-binomial likelihood, two hidden layers and a 100-dimensional latent space, without a batch covariate.

Geneformer ^6^ used the gf-12L-95M-i4096 checkpoint. Tier 1–2 source genes were mapped to human symbols, collapsed when multiple source genes resolved to one identifier, filtered against the model dictionary and rank-tokenized. Geneformer retained 2,297 *D. discoideum* genes and 3,506 *C. elegans* genes. scGPT^7^ used the same Tier 1–2 mappings and retained 2,289 and 3,517 genes, respectively.

Both models produced 512-dimensional cell embeddings.

UCE-4L used the standard pretrained four-layer checkpoint^11^. Inputs used species-specific ESM-2 protein vectors ^27^, chromosome-aware cell sentences and CLS pooling. Protein and chromosome filtering retained 12,548 *D. discoideum* genes and 19,720 *C. elegans* genes. UCE samples expression-weighted gene tokens when constructing cell sentences; coverage therefore denotes the eligible input vocabulary rather than the genes sampled in every cell.

### Coverage and output-dimension sensitivities

The coverage-matched control used the existing UCE-4L embeddings constructed from the Tier 1– 2 ortholog-available source-gene pool; no UCE embedding was rerun for the present analysis. We evaluated this condition for developmental ordering with the same full-data and 30-repeat subsampling protocols as the headline UCE-4L representation.

For the output-dimension control, PCA and scVI were retained unchanged because their headline representations were already 100-dimensional; another 100-component PCA would not reduce their dimensionality. Each frozen scGPT, Geneformer and UCE-4L representation was independently centered and reduced to 100 principal components without feature scaling or whitening, using scikit-learn’s randomized solver and seed 42. The reduction was fit separately for each method and organism and does not place methods in a shared coordinate system. Before writing the full arrays under additive keys, two 512-cell smoke runs produced byte-identical hashes. The reduced arrays were evaluated with the same avgBIO and pseudotime implementations as the headline conditions.

### Ortholog sources, consensus and conflicts

To map source genes into a human vocabulary, we combined OrthoFinder gene trees^52^, OrthoDB eukaryotic orthogroups^53^, OMA pairwise calls ^20^, and a curated organism-specific source (OrthoList 2 for *C. elegans* ^54^; dictyBase for *D. discoideum* ^55^).

Candidate source–human pairs were consolidated by agreement on the same approved NCBI human symbol. Tier 1 required a one-to-one call from both OrthoFinder and OrthoDB plus support from OMA or the curated source. Tier 2 required at least two sources to agree on the same one-to-one target. Tier 3 retained single-source or many-to-many support, and unsupported genes were dropped. When multiple source genes mapped to the same model identifier, expression rows were collapsed before tokenization. The main Geneformer and scGPT conditions used Tiers 1–2; cumulative tier results are reported in Supplementary Table S1.

The number of genes available to a method was counted after its final input filter. PCA and scVI therefore equal the full measured gene set; Geneformer and scGPT include only consensus genes accepted by the respective model vocabulary; UCE includes genes with valid protein and chromosome assets. Percentages use the total measured genes in each AnnData object as denominator.

### Curated annotation construction

We constructed organism-specific annotation categories from published marker resources^34,35,39–41,44^. For each marker set and cell, rank-based enrichment measured the weighted presence of markers among the top 1,500 expressed genes. Null distributions used 50 expression-matched random gene sets, with the 0.99 quantile as the default activity threshold; a two-component Gaussian mixture was used only when it provided a clearly improved and well-supported separation. Empirical null probabilities were adjusted across marker sets within each cell by Benjamini–Hochberg at false-discovery rate 0.05. All six benchmark bundles were assigned as single-label categories: when multiple marker sets were active, the set with the smallest adjusted empirical null probability was selected, and cells with no active set remained unlabeled. These labels are expression-derived annotations rather than independent measurements.

Supplementary Data 1 records the complete curated marker-set memberships, source identifiers, identifier namespaces, panel-level literature context and recorded ortholog tiers. One *C. elegans* marker, opt-2, lacked a stable identifier in the cached symbol mapping; its original symbol is retained and its identifier namespace is listed as unresolved.

### Pseudotime benchmarking

Pseudotime conservation was evaluated with diffusion pseudotime (DPT) ^46^, a graph-based trajectory approach widely used in single-cell benchmarking^56,57^. For each embedding, Scanpy^58^ computed a 15-nearest-neighbor Euclidean graph directly from the representation through use_rep; no shared PCA preprocessing was inserted. The full-data run selected 500 candidate roots uniformly from the earliest developmental state (or all available cells if fewer) and evaluated 50 roots per embedding. DPT was recomputed for each root. Spearman correlation measures rank agreement with the developmental coordinate; its point estimate is the weighted average of the root-specific correlations. The stored consensus pseudotime is the corresponding weighted cell-level average across roots. Consensus Lin’s concordance correlation coefficient (CCC)^47^ is computed once between this consensus pseudotime and the developmental coordinate after pseudotime is mapped to the observed time range, and additionally penalizes scale and location disagreement. Both reported correlations are displayed on [0, 1] by (*r*+1)*/*2.

### Repeated-subsampling stability for pseudotime

Point estimates were computed once on the full data (Table 1). Robustness was then quantified with 30 paired subsampling repeats, each retaining 90% of cells without replacement and drawing 20 roots from the same 500-candidate pool. Every method received the same stratified cell subset and roots in a repeat; its neighbor graph and DPT were recomputed from scratch. Within each repeat, Spearman was averaged across roots and consensus CCC was computed on the root-averaged cell-level pseudotime. Discrete time labels defined strata when there were no more than 20 unique values; otherwise quantile bins of the continuous coordinate were used. We reported the 2.5th–97.5th percentiles across repeats as stability intervals. Because repeats overlap substantially, these are not independent population samples. Pairwise wording refers only to whether the percentile interval of the paired per-repeat difference excludes zero; no null-hypothesis significance test is claimed.

### Biological benchmarking

Annotation preservation followed the scIB framework ^51^. For each embedding and annotation category, Scanpy recomputed a 15-nearest-neighbor Euclidean graph directly from the representation. Leiden clustering was evaluated over 10 candidate resolutions up to 2, retaining the resolution that maximized NMI with the target annotation. We computed NMI^48^, ARI^49^ and full-data label ASW ^50^; avgBIO ^7^ is their unweighted mean. Unknown labels were filtered for the cell-type and milestone bundles and retained for the *D. discoideum* cAMP bundle. Batch-correction metrics were excluded because the endpoint was biological conservation within each atlas. UMAP used 15 neighbors and a minimum distance of 0.1.

### Supporting analyses

To probe gene availability and encoding, we evaluated UCE-4L embeddings restricted to each consensus ortholog tier (Supplementary Table S5), alongside the corresponding tier sweep for the other methods (Supplementary Table S1), and counted retained curated markers (Supplementary Table S2). UCE marker eligibility used the same protein and chromosome filters as the headline UCE condition. These are within-pipeline controls rather than a factorial decomposition of model architecture.

The checkpoint-depth analysis compared the standard pretrained UCE-4L and UCE-33L cell embeddings with the same avgBIO and pseudotime procedures used in the headline evaluation (Supplementary Table S11). Checkpoint sensitivity influenced the choice of UCE-4L for the headline display.

The exploratory protein-space analysis used two previously generated, cell-label-free adapters (Supplementary Table S12). Global moment matching aligned the first two moments of each target-species ESM-2 protein-vector distribution to the human reference distribution. The macrogene-residual adapter began with the global transform and estimated a residual correction from Tier 1 ortholog anchors grouped into protein-space macrogenes. We evaluated mean nearest-human and assigned-ortholog cosine values before and after each transform and evaluated the resulting UCE cell embeddings with the same avgBIO procedure. No cell labels were used to fit either adapter, and these analyses were not used to tune the headline UCE condition.

### Statistical reporting

Gene-availability values are exact counts. Pseudotime bars show the paired stability intervals defined above. avgBIO values are descriptive point estimates. Values are rounded to three decimals in the manuscript.

## Supporting information

Supplementary Data 1: Marker-gene lists

## Data availability

The raw *C. elegans* and *D. discoideum* data are available from GEO under accessions GSE126954 and GSE305468, respectively ^33^. Processed inputs and numerical results are available in the accompanying code repository.

## Code availability

Code for ortholog mapping, embedding generation, evaluation and figure assembly is available at https://github.com/lenatr99/crossspecies-benchmark.

## Supplementary Material

**Figure S1:**
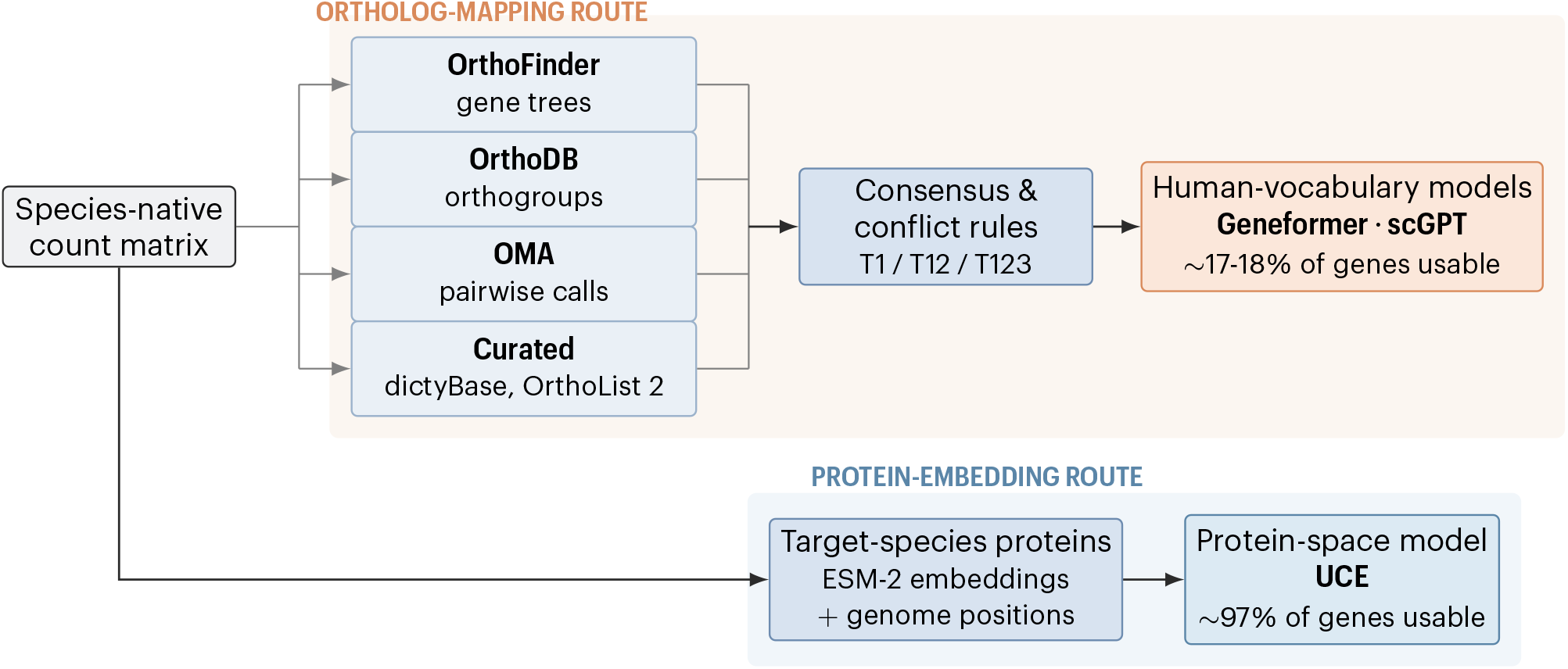
Two input-construction routes from a species-native count matrix. In the ortholog-mapping route, OrthoFinder gene trees, OrthoDB orthogroups, OMA pairwise calls and curated organism resources are combined using consensus and conflict rules before input to the fixed-human-vocabulary Geneformer and scGPT models. In the protein-embedding route, target-species proteins are represented by ESM-2 embeddings and genome positions before input to UCE. Percentages show the proportions of measured genes usable by the corresponding models in the two atlases.

**Table S1:** Ortholog-tier sweep. For each organism, the first row gives the cumulative number of atlas source-gene features selected at Tier 1, Tiers 1–2 and Tiers 1–3 before model-specific identifier and vocabulary filtering. The remaining rows give avgBIO, averaged across the organism’s three annotation panels, for each method restricted to the corresponding tier pool.

| Method / coverage | Tier 1 | Tiers 1–2 | Tiers 1–3 |
|---|---|---|---|
| *D. discoideum* |  |  |  |
| Mapped source genes (recorded input) | 975 | 2337 | 5840 |
| PCA | 0.441 | 0.470 | 0.382 |
| scVI | 0.436 | 0.377 | 0.370 |
| Geneformer | 0.322 | 0.466 | 0.504 |
| scGPT | 0.410 | 0.456 | 0.476 |
| UCE-4L | 0.472 | 0.481 | 0.509 |
| *C. elegans* |  |  |  |
| Mapped source genes (recorded input) | 1659 | 3568 | 9853 |
| PCA | 0.412 | 0.518 | 0.554 |
| scVI | 0.584 | 0.549 | 0.535 |
| Geneformer | 0.371 | 0.549 | 0.596 |
| scGPT | 0.435 | 0.509 | 0.581 |
| UCE-4L | 0.469 | 0.580 | 0.602 |

**Table S2:**
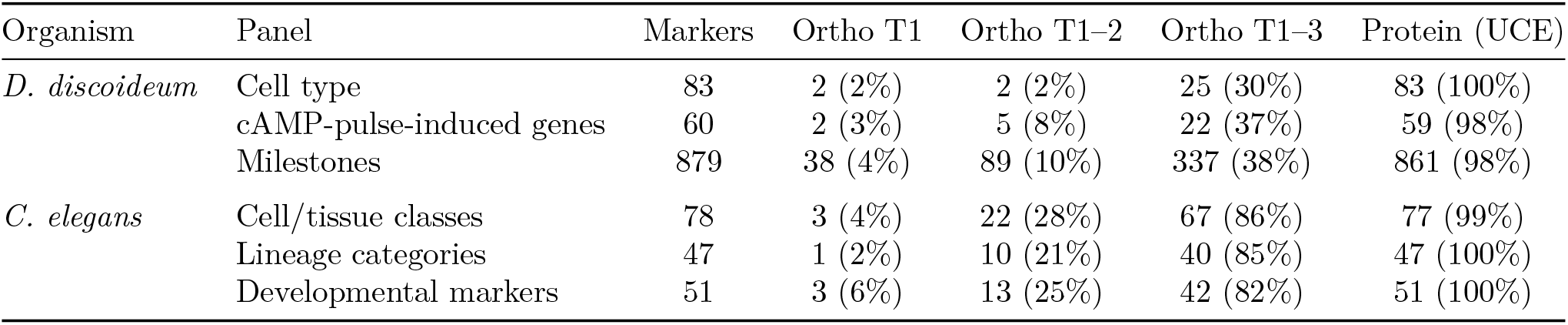
Marker retention across ortholog and protein-based input filters. For each avgBIO annotation panel, the table gives the number of unique marker genes and the number retained at Tier 1, Tiers 1–2 and Tiers 1–3. The UCE column gives eligibility after protein-dictionary and chromosome-position filtering. Tier-pool retention is an upper bound for Geneformer and scGPT because the mapped human ortholog must also occur in each model’s vocabulary. Tier 3 may include single-source or many-to-many support.

**Table S3:**
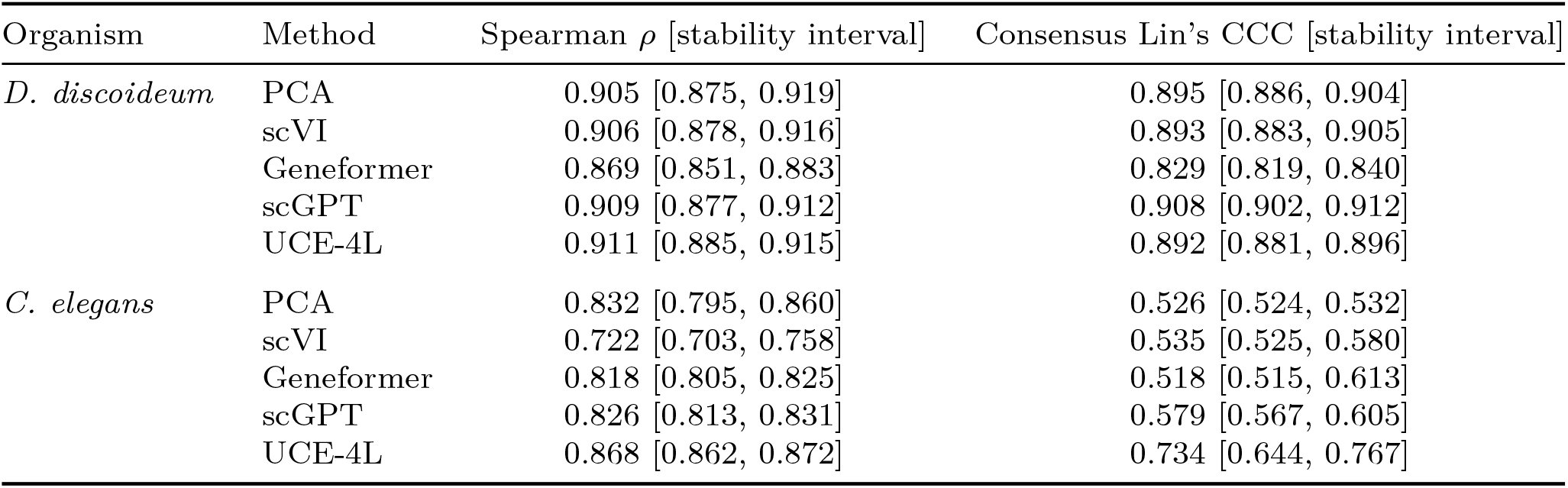
Pseudotime point estimates and repeated-subsampling stability intervals. Point estimates were calculated on the full data. Brackets show the 2.5th–97.5th percentiles across 30 paired repeats, each sampling 90% of cells without replacement and recomputing the neighbour graph and DPT. Spearman is averaged across roots; consensus Lin’s CCC is computed on the root-averaged cell-level pseudotime. All correlations are displayed on [0, 1] via (*r* + 1)*/*2. The brackets are stability intervals.

| Organism | Method | Spearman *ρ* [stability interval] | Consensus Lin’s CCC [stability interval] |
|---|---|---|---|
| *D. discoideum* | PCA | 0.905 [0.875, 0.919] | 0.895 [0.886, 0.904] |
|  | scVI | 0.906 [0.878, 0.916] | 0.893 [0.883, 0.905] |
|  | Geneformer | 0.869 [0.851, 0.883] | 0.829 [0.819, 0.840] |
|  | scGPT | 0.909 [0.877, 0.912] | 0.908 [0.902, 0.912] |
|  | UCE-4L | 0.911 [0.885, 0.915] | 0.892 [0.881, 0.896] |
| *C. elegans* | PCA | 0.832 [0.795, 0.860] | 0.526 [0.524, 0.532] |
|  | scVI | 0.722 [0.703, 0.758] | 0.535 [0.525, 0.580] |
|  | Geneformer | 0.818 [0.805, 0.825] | 0.518 [0.515, 0.613] |
|  | scGPT | 0.826 [0.813, 0.831] | 0.579 [0.567, 0.605] |
|  | UCE-4L | 0.868 [0.862, 0.872] | 0.734 [0.644, 0.767] |

**Table S4:** Component metrics underlying the main avgBIO comparisons. NMI, ARI and label ASW are full-data descriptive point estimates; avgBIO is their unweighted mean. Geneformer and scGPT use Tier 1–2 ortholog mappings, and UCE-4L denotes the standard pretrained four-layer model.

| Organism | Annotation | Method | NMI | ARI | ASW | avgBIO |
|---|---|---|---|---|---|---|
| *D. discoideum* | Cell type | PCA | .393 | .359 | .558 | .437 |
|  |  | scVI | .362 | .258 | .583 | .401 |
|  |  | Geneformer | .412 | .410 | .549 | .457 |
|  |  | scGPT | .224 | .193 | .550 | .322 |
|  |  | UCE-4L | .590 | .724 | .539 | .618 |
|  | cAMP-pulse-induced genes | PCA | .450 | .279 | .644 | .458 |
|  |  | scVI | .357 | .148 | .581 | .362 |
|  |  | Geneformer | .621 | .618 | .626 | .622 |
|  |  | scGPT | .794 | .884 | .584 | .754 |
|  |  | UCE-4L | .799 | .890 | .654 | .781 |
|  | Milestones | PCA | .306 | .239 | .416 | .321 |
|  |  | scVI | .279 | .156 | .514 | .316 |
|  |  | Geneformer | .264 | .194 | .501 | .319 |
|  |  | scGPT | .206 | .184 | .490 | .293 |
|  |  | UCE-4L | .230 | .170 | .494 | .298 |
| *C. elegans* | Cell/tissue classes | PCA | .739 | .775 | .457 | .657 |
|  |  | scVI | .721 | .601 | .576 | .633 |
|  |  | Geneformer | .559 | .636 | .533 | .576 |
|  |  | scGPT | .575 | .633 | .538 | .582 |
|  |  | UCE-4L | .725 | .821 | .550 | .699 |
|  | Lineage categories | PCA | .620 | .606 | .482 | .569 |
|  |  | scVI | .529 | .299 | .567 | .465 |
|  |  | Geneformer | .598 | .606 | .543 | .582 |
|  |  | scGPT | .567 | .543 | .551 | .554 |
|  |  | UCE-4L | .633 | .637 | .562 | .610 |
|  | Developmental markers | PCA | .410 | .467 | .526 | .468 |
|  |  | scVI | .347 | .150 | .523 | .340 |
|  |  | Geneformer | .460 | .496 | .511 | .489 |
|  |  | scGPT | .355 | .313 | .510 | .393 |
|  |  | UCE-4L | .494 | .516 | .518 | .509 |

**Table S5:** Ortholog-pool-restricted UCE comparison. avgBIO for full UCE-4L, UCE-4L restricted to the Tier 1–2 source-gene pool used before each model’s vocabulary filter, Geneformer and scGPT. The best value in each row is shown in bold.

| Organism | Panel | UCE-4L (full) | UCE-4L (Tier 1–2) | Geneformer | scGPT |
|---|---|---|---|---|---|
| *D. discoideum* | Cell type | **0.618** | 0.342 | 0.457 | 0.322 |
|  | cAMP-pulse-induced genes | **0.781** | 0.770 | 0.622 | 0.754 |
|  | Milestones | 0.298 | **0.332** | 0.319 | 0.293 |
| *C. elegans* | Cell/tissue classes | **0.699** | 0.655 | 0.576 | 0.582 |
|  | Lineage categories | **0.610** | 0.593 | 0.582 | 0.554 |
|  | Developmental markers | **0.509** | 0.491 | 0.489 | 0.393 |

**Table S6:** Coverage-matched UCE-4L sensitivity for developmental ordering. The full UCE-4L representation uses all eligible target-species proteins; the matched condition is reconstructed from genes with Tier 1–2 source-to-human ortholog assignments. Values are full-data point estimates with 2.5th–97.5th percentile intervals from 30 repeated 90% subsamples. Correlations are displayed on [0, 1] via (*r* + 1)*/*2.

| Organism | UCE-4L gene set | Spearman *ρ* [interval] | Consensus Lin's CCC [interval] |
|---|---|---|---|
| *D. discoideum* | Full eligible gene set | 0.911 [0.885, 0.915] | 0.892 [0.881, 0.896] |
|  | Ortholog-available gene set | 0.914 [0.889, 0.915] | 0.903 [0.893, 0.909] |
| *C. elegans* | Full eligible gene set | 0.868 [0.862, 0.872] | 0.734 [0.644, 0.767] |
|  | Ortholog-available gene set | 0.868 [0.863, 0.871] | 0.708 [0.589, 0.763] |

**Table S7:** PCA preprocessing sensitivity for annotation preservation. Raw-count PCA is the headline condition. The preprocessed condition selects 2,000 Seurat-v3 highly variable genes, normalizes to 10,000 counts per cell, applies log1p, scales with a cap of 10 and computes 100 PCs. Delta is preprocessed minus raw.

| Organism | Annotation | Raw PCA | Preprocessed PCA | Delta |
|---|---|---|---|---|
| *D. discoideum* | Cell type | 0.437 | 0.531 | 0.094 |
|  | cAMP-pulse-induced genes | 0.458 | 0.484 | 0.026 |
|  | Milestones | 0.321 | 0.387 | 0.066 |
| *C. elegans* | Cell/tissue classes | 0.657 | 0.724 | 0.067 |
|  | Lineage categories | 0.569 | 0.643 | 0.073 |
|  | Developmental markers | 0.468 | 0.387 | -0.080 |

**Table S8:**
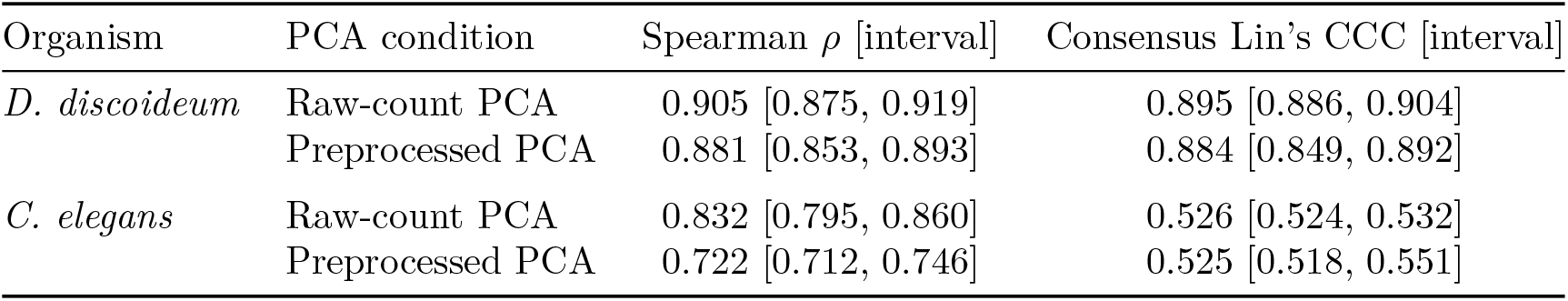
PCA preprocessing sensitivity for developmental ordering. Full-data point estimates and 2.5th–97.5th percentile stability intervals from 30 repeated 90% subsamples; correlations are displayed on [0, 1] via (*r* + 1)*/*2.

| Organism | PCA condition | Spearman *ρ* [interval] | Consensus Lin's CCC [interval] |
|---|---|---|---|
| *D. discoideum* | Raw-count PCA | 0.905 [0.875, 0.919] | 0.895 [0.886, 0.904] |
|  | Preprocessed PCA | 0.881 [0.853, 0.893] | 0.884 [0.849, 0.892] |
| *C. elegans* | Raw-count PCA | 0.832 [0.795, 0.860] | 0.526 [0.524, 0.532] |
|  | Preprocessed PCA | 0.722 [0.712, 0.746] | 0.525 [0.518, 0.551] |

**Table S9:** Common 100-component sensitivity for annotation preservation. PCA and scVI are retained at their native 100 dimensions; scGPT, Geneformer and UCE-4L are independently reduced to 100 components by centered, unscaled PCA. Cells show native *→* 100-dimensional avgBIO, with the change in parentheses. The lower block gives the fraction of variance retained by the 100 components.

| Organism | Annotation | PCA | scVI | scGPT | Geneformer | UCE-4L |
|---|---|---|---|---|---|---|
| *D. discoideum* | Cell type | 0.437 → 0.437 (+0.000) | 0.401 → 0.401 (+0.000) | 0.322 → 0.353 (+0.031) | 0.457 → 0.457 (+0.000) | 0.618 → 0.607 (-0.011) |
|  | cAMP-pulse-induced genes | 0.458 → 0.458 (+0.000) | 0.362 → 0.362 (+0.000) | 0.754 → 0.753 (-0.001) | 0.622 → 0.639 (+0.017) | 0.781 → 0.782 (+0.001) |
|  | Milestones | 0.321 → 0.321 (+0.000) | 0.316 → 0.316 (+0.000) | 0.293 → 0.280 (-0.014) | 0.319 → 0.321 (+0.002) | 0.298 → 0.311 (+0.013) |
| *C. elegans* | Cell/tissue classes | 0.657 → 0.657 (+0.000) | 0.633 → 0.633 (+0.000) | 0.582 → 0.573 (-0.009) | 0.576 → 0.592 (+0.016) | 0.699 → 0.698 (+0.000) |
|  | Lineage categories | 0.569 → 0.569 (+0.000) | 0.465 → 0.465 (+0.000) | 0.554 → 0.558 (+0.005) | 0.582 → 0.580 (-0.003) | 0.610 → 0.612 (+0.002) |
|  | Developmental markers | 0.468 → 0.468 (+0.000) | 0.340 → 0.340 (+0.000) | 0.393 → 0.394 (+0.002) | 0.489 → 0.489 (+0.000) | 0.509 → 0.510 (+0.001) |
| Organism |  | PCA | scVI | scGPT | Geneformer | UCE-4L |
| *D. discoideum* |  | 1.000 | 1.000 | 0.973 | 0.960 | 0.981 |
| *C. elegans* |  | 1.000 | 1.000 | 0.951 | 0.953 | 0.969 |

**Table S10:**
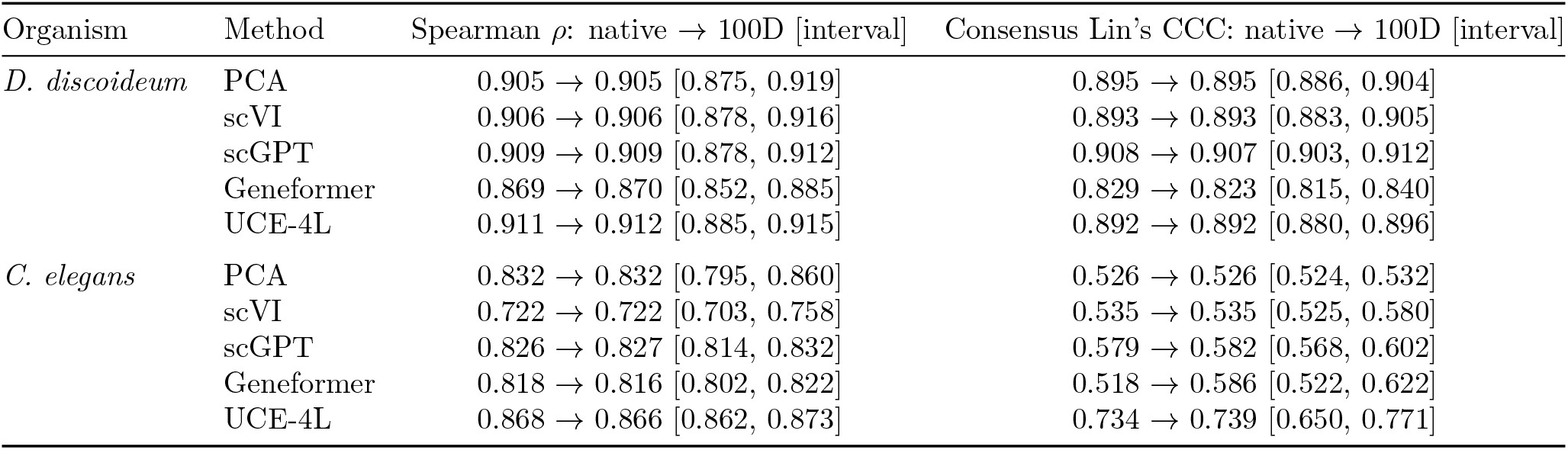
Common 100-component sensitivity for developmental ordering. PCA and scVI are retained at their native 100 dimensions; scGPT, Geneformer and UCE-4L are reduced by post-embedding PCA. Each cell shows the native and 100-dimensional full-data point estimates, followed by the 2.5th–97.5th percentile stability interval for the 100-dimensional condition from 30 repeated 90% subsamples. Correlations are displayed on [0, 1] via (*r* + 1)*/*2.

| Organism | Method | Spearman *ρ*: native → 100D [interval] | Consensus Lin's CCC: native → 100D [interval] |
|---|---|---|---|
| *D. discoideum* | PCA | 0.905 → 0.905 [0.875, 0.919] | 0.895 → 0.895 [0.886, 0.904] |
|  | scVI | 0.906 → 0.906 [0.878, 0.916] | 0.893 → 0.893 [0.883, 0.905] |
|  | scGPT | 0.909 → 0.909 [0.878, 0.912] | 0.908 → 0.907 [0.903, 0.912] |
|  | Geneformer | 0.869 → 0.870 [0.852, 0.885] | 0.829 → 0.823 [0.815, 0.840] |
|  | UCE-4L | 0.911 → 0.912 [0.885, 0.915] | 0.892 → 0.892 [0.880, 0.896] |
| *C. elegans* | PCA | 0.832 → 0.832 [0.795, 0.860] | 0.526 → 0.526 [0.524, 0.532] |
|  | scVI | 0.722 → 0.722 [0.703, 0.758] | 0.535 → 0.535 [0.525, 0.580] |
|  | scGPT | 0.826 → 0.827 [0.814, 0.832] | 0.579 → 0.582 [0.568, 0.602] |
|  | Geneformer | 0.818 → 0.816 [0.802, 0.822] | 0.518 → 0.586 [0.522, 0.622] |
|  | UCE-4L | 0.868 → 0.866 [0.862, 0.873] | 0.734 → 0.739 [0.650, 0.771] |

**Table S11:** UCE checkpoint-depth sensitivity. The upper block reports full-data avgBIO point estimates. The lower block reports full-data developmental-ordering estimates with 2.5th–97.5th percentile stability intervals from 30 repeated 90% subsamples. Correlations are displayed on [0, 1] via (*r* + 1)*/*2.

| Organism | Annotation | UCE-4L | UCE-33L |
|---|---|---|---|
| *D. discoideum* | Cell type | 0.618 | 0.252 |
|  | cAMP-pulse-induced genes | 0.781 | 0.500 |
|  | Milestones | 0.298 | 0.250 |
| *C. elegans* | Cell/tissue classes | 0.699 | 0.717 |
|  | Lineage categories | 0.610 | 0.614 |
|  | Developmental markers | 0.509 | 0.514 |
| Organism | Metric | UCE-4L [interval] | UCE-33L [interval] |
| *D. discoideum* | Spearman *ρ* | 0.911 [0.885, 0.915] | 0.704 [0.683, 0.719] |
|  | Consensus Lin's CCC | 0.892 [0.881, 0.896] | 0.904 [0.819, 0.920] |
| *C. elegans* | Spearman *ρ* | 0.868 [0.862, 0.872] | 0.851 [0.846, 0.855] |
|  | Consensus Lin's CCC | 0.734 [0.644, 0.767] | 0.646 [0.611, 0.679] |

**Table S12:** UCE protein-space calibration sensitivity. The upper block reports full-data avgBIO before and after two cell-label-free protein-space adapters. “Global” denotes global moment matching; “macrogene” denotes global matching followed by a residual transform estimated from Tier 1 anchor pairs. The lower block reports mean nearest-human and assigned-ortholog cosine values for the same raw and adapted protein vectors. Global matching used 13,209 *D. discoideum* or 19,985 *C. elegans* target proteins and one shared set of 19,790 human proteins; the macrogene adapter used 960 or 1,638 Tier 1 anchor pairs, respectively.

| Organism | Annotation | 4L raw | 4L global | 33L raw | 33L global | 33L macrogene |
|---|---|---|---|---|---|---|
| *D. discoideum* | Cell type | 0.618 | 0.456 | 0.252 | 0.430 | 0.591 |
|  | cAMP-pulse-induced genes | 0.781 | 0.778 | 0.500 | 0.771 | 0.769 |
|  | Milestones | 0.298 | 0.329 | 0.250 | 0.306 | 0.303 |
| *C. elegans* | Cell/tissue classes | 0.699 | 0.692 | 0.717 | 0.689 | 0.661 |
|  | Lineage categories | 0.610 | 0.612 | 0.614 | 0.607 | 0.603 |
|  | Developmental markers | 0.509 | 0.508 | 0.514 | 0.508 | 0.510 |
| Organism | Protein diagnostic | Raw | Global | Macrogene | Evaluation *n* |  |
| *D. discoideum* | Nearest human | 0.761 | 0.806 | 0.809 | 13,209 |  |
|  | Assigned ortholog T1 | 0.721 | 0.810 | 0.815 | 960 |  |
|  | Assigned ortholog T2 | 0.731 | 0.807 | 0.812 | 1,331 |  |
|  | Assigned ortholog T3 | 0.661 | 0.764 | 0.767 | 3,415 |  |
| *C. elegans* | Nearest human | 0.813 | 0.828 | 0.829 | 19,985 |  |
|  | Assigned ortholog T1 | 0.781 | 0.832 | 0.835 | 1,638 |  |
|  | Assigned ortholog T2 | 0.801 | 0.835 | 0.838 | 1,809 |  |
|  | Assigned ortholog T3 | 0.730 | 0.784 | 0.786 | 6,085 |  |

## Notes

https://github.com/lenatr99/crossspecies-benchmark

https://www.ncbi.nlm.nih.gov/geo/query/acc.cgi?acc=GSE126954

https://www.ncbi.nlm.nih.gov/geo/query/acc.cgi?acc=GSE305468

### Competing Interest Statement

The authors have declared no competing interest.

